# The role of IL-6 in dopamine dysregulation underlying anhedonia phenotype in rats

**DOI:** 10.1101/2023.11.21.568169

**Authors:** Roger B Varela, Heather Macpherson, Tristan Houghton, Dara Daygon, Susannah J Tye

## Abstract

**Aims:** To investigate the role of peripheral metabolic change and chronic low-grade inflammation on striatal dopamine dynamics and anhedonia-like behaviour induced by hypothalamic–pituitary–adrenal (HPA) axis disruption.

**Methods:** Wistar rats were trained in a progressive-ratio/concurrent effort-related choice paradigm to assess effort-related decision making. After reaching a stable baseline, animals received daily injections of adrenocorticotrophic hormone (ACTH) or saline for 24 days. On the 23^rd^ and 24^th^ day, animals received a bupropion challenge (10mg/kg and 20mg/kg respectively) 30 minutes prior to the behavioural testing session. On the 25^th^ day, animals received a single injection of bupropion (20mg/kg) 30 minutes prior to euthanasia. Peripheral and central inflammatory markers were assessed through ELISA and In-Cell Western assay; glucose transport activity was assessed in peripheral blood mononuclear cells though a commercial assay kit; brain levels of dopaminergic and inflammatory markers were assessed in the nucleus accumbens (NAc) and prefrontal cortex (PFC) through immunohistochemistry; and serum central carbon metabolism metabolites were assessed through a metabolomics approach.

**Results:** ACTH induced an anhedonia-like phenotype, decreased tyrosine hydroxylase (TH) levels in the NAc, increased peripheral IL-6 levels, and decreased glucose transport activity and glucose metabolites when compared to control group. Bupropion treatment was not able to reverse the anhedonic phenotype. Glucose uptake was positively correlated to behaviour; TH levels were correlated to microglia volume; metabolites were correlated to TH levels; and IL6 was correlated to TH levels and metabolites.

**Conclusion:** Chronic ACTH treatment can induce treatment-resistant anhedonia in rats, and the interaction between peripheral immunometabolic state and central dopamine synthesis is a potential mechanism underlying this phenotype.

## INTRODUCTION

Dopaminergic dysfunction is the hallmark for the pathophysiology of several psychiatric disorders including depression, addiction, bipolar disorder and schizophrenia. Dopamine (DA) is a neurotransmitter produced by neurons in the midbrain, which projects to several regions in the brain, including striatal, limbic and cortical areas [1]. The mesolimbic pathway consists of DAergic projections from the ventral tegmental area (VTA) to the nucleus accumbens (NAc), and is known to regulate reward-related positive and negative reinforcement, incentive salience, aversion-related cognition and decision-making [2]. It is also considered to be the central hub for control of motivation [3]. Activation of DA neurons in the VTA increases NAc activity, which is correlated to reward-seeking behaviour in rodents [4]. In humans, activation of the NAc is observed in response to anticipation of reward during monetary behaviour tasks, and this activation is reduced in subjects with depressive symptoms, especially anhedonia [5].

According to DSM-5, anhedonia is defined by either a reduced ability to experience pleasure or a diminished interest in engaging in pleasurable activities, also known as motivational anhedonia [6]. Anhedonia is a multi-faceted symptom observed in several psychiatric disorders, being a core feature of depression [7]. The hypothesis of DAergic dysfunction underlying the neurobiology of anhedonia symptoms is well known. The mesolimbic system is affected in mood disorder patients, and a reduction of DA transmission in the NAc leads to the development of anhedonia symptoms, as shown by neuroimaging and pharmacological manipulation studies [7]. Although its precise biological triggers are not fully understood, some evidence suggests that inflammation may be related to DA dysfunction and anhedonia. According to Miller et al. [8] the activation of the inflammasome induced by stress and non-pathogenic stimuli may drive a peripheral inflammatory response, which can traffic to the brain impacting several neurotransmitter systems, including DA. This relationship between inflammatory pathways and the brain may drive depression and contribute to non-response to antidepressant medication.

Around 30% of patients with major depressive disorder do not respond adequately to classical antidepressants such as citalopram, imipramine and the DA modulator bupropion, classifying treatment resistant depression (TRD) [9, 10]. Evidence suggests the importance of anhedonia and peripheral inflammation as predictive biomarkers to antidepressant response. Depressed patients have increased levels of peripheral markers of inflammation such IL-6 [11, 12]. IL-6 levels are also higher in depressive patients with than without anhedonia, where it is correlated to severity of symptoms [13]. C-reactive protein (CRP), another peripheral marker of inflammation, is also associated with decreased functional connectivity in the reward system [14], supporting the relation between inflammation and anhedonia. By itself, baseline levels of anhedonia are a negative prognostic factor for depression and can predict poorer recovery, longer time to respond and resistance to classical antidepressants [15] and transcranial magnetic stimulation [16]. In addition, anhedonia represents a major risk factor for suicide [17]. Interestingly, some studies have demonstrated that anhedonia is a clinical predictor for effectiveness of therapies used for treatment-resistant-depression, such as ketamine [18].

Translational animal models are relevant tools to investigate the biological mechanisms and circuits underlying normal and pathological behaviours, as well as responsiveness to different treatment approaches [19]. According to the Research Domain Criteria (RDoC) system, psychiatric symptoms can be categorised into five main behavioural domains and endophenotype. These domains are based on well-known evolutionary, anatomical, and network properties, and thus are the most appropriate approach for translational research [20]. RDoC’s positive valence behaviour domain relates to the endophenotype which includes responsiveness to reward, reward learning, habits, approach, and motivation. Since mesolimbic circuitry anatomy is conserved throughout mammalian evolution, and reward-related behaviours can be translationally assessed [21], rodent models are valid tools to investigate the link between chronic stress, inflammation and effort related decision making deficits.

Chronic administration of adrenocorticotropic hormone (ACTH) is known for inducing a classical antidepressant-resistant state in rats, and inflammation and DA dysregulation seem to play key roles in this phenotype [22, 23]. However, the effects of ACTH on mesolimbic DA and anhedonia-like phenotype are not fully characterised yet. The present study aims to describe the effects of chronic ACTH on 1) anhedonia-like phenotype induced using the effort-related choice test; 2) associated mesolimbic DAergic alteration; 3) peripheral and central inflammatory alterations; and, 4) peripheral glucose metabolism disturbances.

## METHODS

### Animals

The subjects were adult male Wistar rats (N=16) obtained from the Animal Resources Centre, transported, and housed at the University of Queensland animal facility. The animals were paired housed with a divider and maintained on a 12-h light/dark cycle (lights on at 6:00 a.m.) at a temperature of 22±1 °C. All experimental procedures were performed in accordance with, and with the approval of the University of Queensland Institutional Animal Care and Use Committees (QBI/267/20). All experiments were performed at the same time during the day to mitigate circadian variations.

### Drugs

The drugs used in this study included the following: human ACTH 1-24 (AnaSpec, USA), 150μg per animal dissolved in 0.9% saline; bupropion hydrochloride (Merck, Germany), 10 or 20mg per animal per day dissolved in 0.9% saline; control vehicle 0.9% saline; and Lethabarb (Virbac Animal Health, Australia; constituents: 325mg/ml sodium pentobarbitone and 100μg/ml buprenorphine), 1ml per animal. Drugs were administered via intraperitoneal injection.

### Experimental design

#### Anhedonia-like phenotype – Effort related choice task (ERCT)

The ERCT utilised in this experiment was adapted from Randall and colleagues [24]. Animals were initially food restricted to 80% of the body weight of free-feed animals. The behavioural sessions (30-minute sessions, five days a week) were conducted in operant conditioning chambers (30.5cm x 25.4cm x 30.5cm; Coulbourn Instruments, USA) during the light period. Rats were initially trained to press a lever on a fixed-ratio schedule (FR1; one lever press for one pellet) to obtain 45 mg dustless sugar pellets (Able Scientific, AUS) for a week before being shifted to the PROG schedule. During the PROG sessions, the ratio requirement started at FR1 and increased by one additional response every time 15 pellets were dispensed. Once the animals reached a stable number of lever presses over four days (less than 20% difference), the PROG/CONC paradigm started. In this paradigm, 10g of laboratory chow was made concurrently available in the chamber during each PROG session, so that the animals had a choice between approaching and consuming chow (low-effort, low-value) or pressing a lever on a PROG schedule for a sucrose pellet (high-effort, high-value). After reaching a stable baseline in the PROG/CONC paradigm (pre-treatment), animals were separated into control (n=8) or ACTH (n=8) groups (so that each group had an equal number of low and high responders) and received daily injections of saline or ACTH for 24 days while training sessions were kept as usual. Finally, all animals received injections of 10mg bupropion hydrochloride on day 23 of treatment and 20mg bupropion hydrochloride on day 24, thirty minutes prior to behavioural testing. At the end of each session, food intake was determined by weighing the remaining chow, including spillage. Number of rewards, lever presses and highest ratio achieved were also recorded with Graphic State software (Coulbourn Instruments, USA). Rats were euthanised with Lethabarb thirty minutes after intraperitoneal injection of 20mg bupropion hydrochloride, one day after completion of behavioural testing. Blood was taken via cardiac puncture, serum was isolated through centrifugation and peripheral blood mononuclear cells (PBMCs) were isolated through Lymphoprep™ density gradient medium according to manufacturer instructions (Stemcell Technologies, Germany). The brains were then removed, left hemisphere was rapid frozen in liquid nitrogen and stored at -80C, and right hemisphere fixed in 4% paraformaldehyde (24hr), then immersed in 0.1% sodium azide in phosphate-buffered saline (PBS) solution for posterior immunohistochemistry (IHC) analysis.

#### Central inflammatory and DAergic markers – Immunohistochemistry and ELISA

40μm thickness sections including NAc core (NAcC), NAc shell (NAcS), prelimbic cortex (PrLC) and infralimbic cortex (ILC) regions were obtained using a sliding microtome (Leica Microsystems, Germany). Sections were incubated in antigen recovery solution (0.1M sodium citrate, citric acid added until pH reaches 6, 0.1M SDS) at 40°C for thirty minutes using a shaker incubator. After antigen recovery, the samples were washed in water and PBS before being incubated in blocking solution (0.5% bovine serum albumin in 0.05% saponin, 0.05% sodium azide in 0.1M PBS) for thirty minutes. The tissue was then incubated for at least 72 hours in primary antibodies (prepared in blocking solution at 4ºC). Primary antibodies were then removed, and tissue sections were washed thrice in PBS before incubating them in blocking solution for five minutes, and then in appropriate species-specific biotinylated secondary antibodies overnight. The primary antibodies used for immunohistochemical analysis included: Anti-tyrosine hydroxylase (mouse monoclonal, 1:2000, 22941, ImmunoStar, USA), anti-neuronal nuclear protein (NeuN) (rabbit monoclonal, 1:2000, 24307S, Cell Signalling Technology, USA), anti-ionised calcium binding adaptor molecule (IBA1) (Rabbit monoclonal, 1:1000, 016-26721, FUJIFILM Wako, Japan). Secondary antibodies included: goat anti-mouse IgG Alexa Fluor 555 (1:1000, A-21424, Thermo Fisher, USA), goat anti-rabbit IgG Alexa Fluor 488 (1:1000, A-11034, Thermo Fisher, USA), Hoescht 33342 (1:2000, H3570, Thermo Fisher, USA). After incubation, the tissue sections were washed thrice in PBS, mounted on adhesive-coated, positively charged slides (InstrumeC, AUS) with fluorescent mounting medium (Agilent Technologies, USA) and sealed with nail polish. All tissue imaging was conducted using the Axio Imager Azure microscope (ZEISS©, Germany) using a single field of view from the structure of interest, captured using a 20x objective lens for DAergic and neural markers and 40x for microglial markers. LED exposure and light settings were kept consistent for all sections belonging to a group. The images were then analysed with semi-quantitative immunofluorescent intensity analysis using Fiji (ImageJ) [25]. Identical threshold settings were used between images to ensure consistency.

Brain IL-6 levels were determined using a commercially available ELISA kit (RayBiotech, USA). First, 100mg brain sections were collected from frozen tissue (AP:0.9 from bregma) and lysed in 500 μL cold radioimmunoprecipitation assay (RIPA) buffer. Tissue lysis was performed by sonication for 10-20 seconds followed by centrifugation (1600 rpm, 20 minutes at 4 °C). The supernatant was collected and kept at -80ºC until further analysis. The ELISA was performed in brain extracts according to the manufacturer’s instructions.

#### Peripheral inflammatory markers – ELISA and In-Cell Western assay

Serum concentrations of CRP and IL-6 were determined using a commercially available ELISA kit (Thermo Fisher, USA) according to the manufacturer’s instructions. For NF-κB levels, after isolation PBMCs were seeded in a 96-well plate at a density of 50,000 cells per well, incubated for thirty minutes at room temperature, then in ice-cold methanol for twenty minutes. Cells were washed thrice with PBS with 0.2% Tween-20 then blocked in Intercept Blocking Buffer (Li-Cor, USA) for 90 minutes. PBMCs were then incubated in a primary anti-NF-κB (Rabbit monoclonal, 1:200; Cell Signalling Technology, USA) antibody overnight at 4ºC under gentle agitation. Cells were washed in PBS four times and then incubated in the secondary antibody goat anti-rabbit IgG IRDye 800CW (1:1000; Li-Cor, USA) and Cell Tag 700 Stain (1:500, Li-Cor, USA) for one hour at room temperature under gentle agitation. Finally, cells were washed four times in PBS. Cell Tag and NF-κB fluorescence were scanned with the Odyssey CLx Imaging system (Li-Cor, USA) at 169μM.

#### Peripheral metabolic markers – Glucose uptake commercial kit and metabolomic approach

PBMC glucose uptake was determined using Glucose Uptake Cell-based Assay kit^®^ (Cayman chemical, USA) according to manufacturer’s instructions, after five minute incubation with insulin or growth media only. No treatment was performed to seeded cells, except an insulin or vehicle challenge (in duplicates) five minutes before the end of overnight incubation with fluorescently labelled deoxyglucose analog (2-NBDG). Fluorescence from 2-NBDG taken up by the cells was scanned with the Odyssey CLx Imaging system.

Central carbon metabolism (CCM) markers were analysed through liquid chromatography–mass spectrometry (LC/MS) in serum samples. Briefly, 100μl of serum sample was mixed with extraction buffer (50% MeOH, 1ml/100ml 500μM AZT, 0.5% AA) for metabolite extraction. 100μl of phenol:chloroform:Isoamyl-alcohol was added to the samples and centrifuged for 16000RCF for 5 minutes at 4ºC. Supernatant was transferred to new tubes and evaporated for 30 minutes in a vacuum concentrator. 200μl of water was added to the tubes;the extracts were then freeze-dried overnight and reconstituted in 200ul 2%ACN (twice the volume of the original sample volume used). The samples were injected in the LC at 2μl and 10μl to be able to capture both low and high concentration metabolites. Metabolite concentrations were corrected by dilution factor. The full panel of metabolites assessed in the present study is described on supplementary table 1.

### Statistical analysis

All statistical analyses were performed on GraphPad Prism 9.1.2 (GraphPad Software Inc., USA). Normality of data were determined using Shapiro-Wilk and D’Agostino-Pearson tests, and variance by calculating F-ratios. Mean differences between groups were examined using unpaired two-tailed t-tests; repeated-measures one-way ANOVAs with Dunnett’s post hoc analyses on normal data, and Mann-Whitney two-tailed U-tests or Friedman tests with Dunn’s post hoc analyses on non-normal data; or two-way ANOVA with Tukey’s post hoc analysis. Welch’s or Geisser-Greenhouse corrections were applied to results from groups with unequal variance. Correlations analysis was performed with Pearson’s test on normal data and Spearman’s test on non-normal data. Relationships that were found to be significant and relevant were selected and tested using linear regression, followed by Holm-Šídák post hoc analyses for multiple comparisons. A p-value of less than 0.05 was considered statistically significant for all results.

## RESULTS

Chronic ACTH injections for 25 days were able to decrease the number of lever presses [F(3,7)=15.7, p=0.0013], number of rewards [F(3,7)=21.2, p<0.0001], highest ratio achieved [F(3,7)=12.0, p=0.0011] and chow consumption [F(3,7)=25.5, p<0.0001] in the ERCT when compared to pre-treatment period from the same animals (fig 1A-D). Acute administration of either 10mg or 20mg bupropion was not able to reverse ACTH-induced suppression of effort-related choice behaviour. The overall treatment effect of daily saline injections was statistically significant for chow consumption [F(3,7)=23.1, p=0.0002] but not for lever pressing, number of rewards, or highest ratio achieved (Fig 1E-H). Furthermore, acute administration of either 10mg or 20mg bupropion had no effect on these behavioural measures in saline-treated animals. Unpaired t-tests between experimental groups revealed that chronic ACTH administration significantly decreased lever pressing (t=2.26, difference between means ± SEM = 0.319 ± 0.141, p=0.0417) and number of rewards (t=2.25, difference between means ± SEM = 0.259 ± 0.115, p=0.0426) compared to control treatment (Fig 1I-L). There were no significant differences in highest ratio achieved or chow consumption between ACTH and saline-treated control animals. Together these results demonstrate that chronic ACTH injection was able to induce an anhedonia-like phenotype in rats.

**Figure 1:**
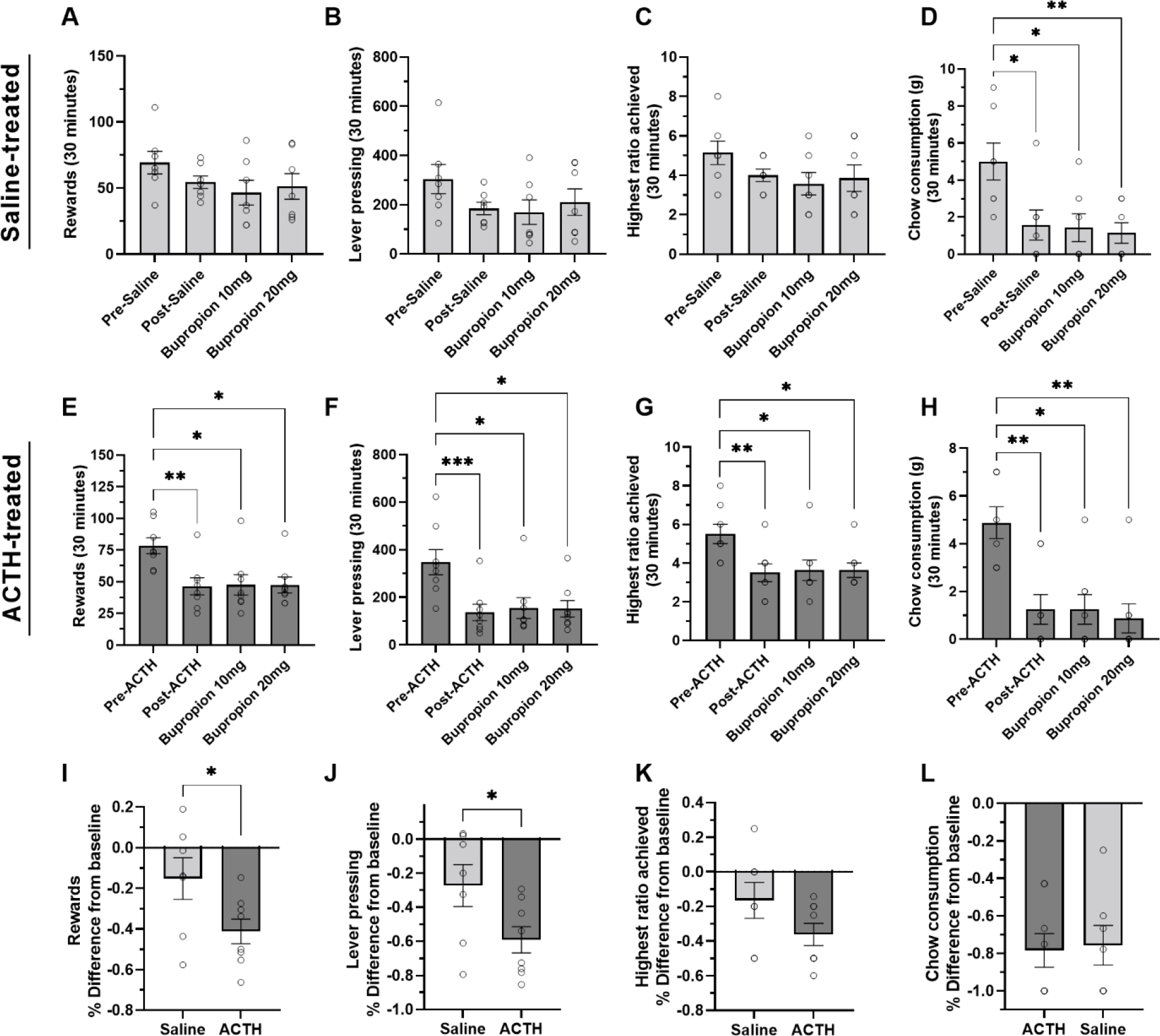
The effects of chronic adrenocorticotrophin (ACTH) or saline administration on effort-related choice behaviour. Mean (±SEM) number of lever presses **(A, E)**; number of rewards **(B**,**F)**; highest ratio achieved **(C**,**G)**; and chow consumption **(D, H)** during 30-minute effort-related choice task sessions by ACTH-treated animals compared between pre-treatment, post-treatment, bupropion 10mg injection and bupropion 20mg injection stages. *P<0.05; **P<0.01; ***P<0.001 according to repeated measures one-way analysis of variance with Geisser-Greenhouse correction, followed by Dunnett’s post hoc test. Percentage difference in lever pressing **(I)**; number of rewards **(J)**; highest ratio achieved **(K)**; and chow consumption **(L)** between baseline and the final effort-related choice task session after chronic treatment with ACTH or saline pre-bupropion. *p<0.05; ns = non-significant, according to unpaired t-test.

Unpaired t-tests between experimental groups revealed chronic ACTH administration significantly decreased TH immunofluorescence in the NAcS (U=6, p=0.0093) and ILC (t=2.494, p=0.0269) compared to saline treatment (Fig 2).

**Figure 2:**
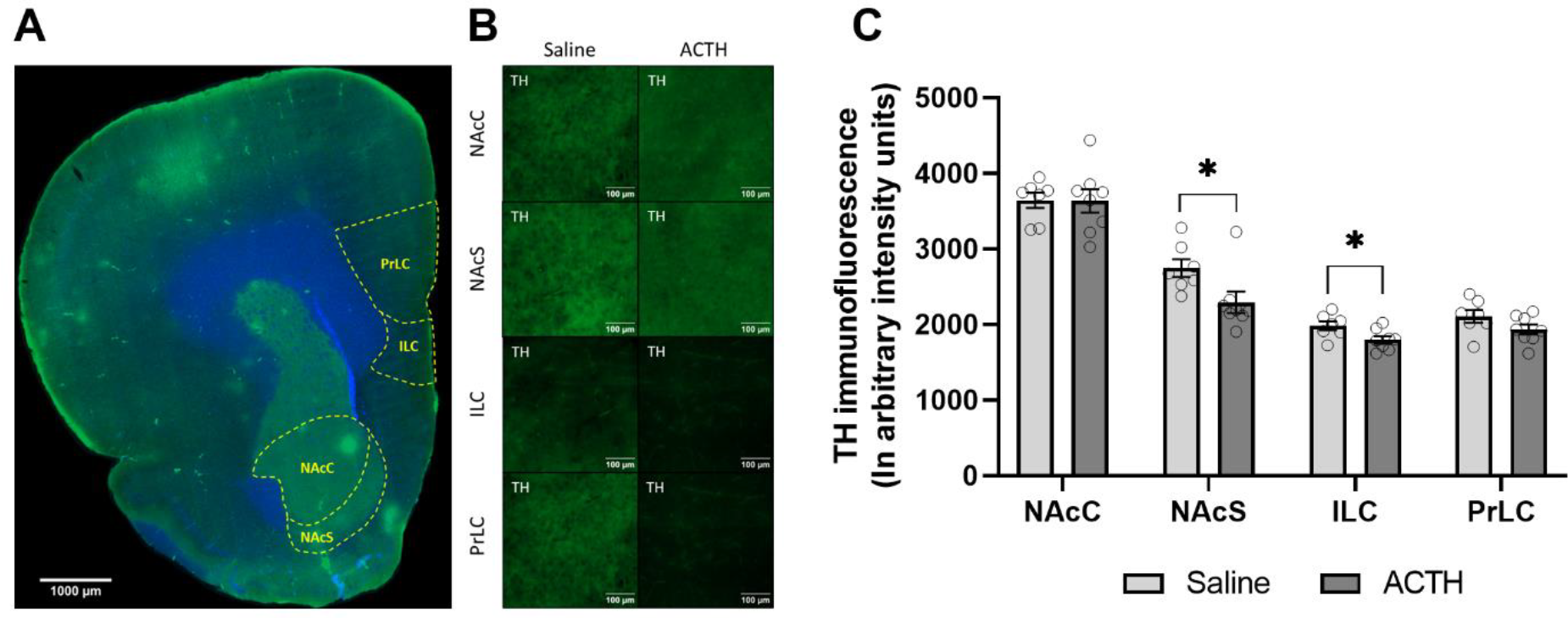
Effects of chronic adrenocorticotrophic hormone (ACTH) or saline treatment on tyrosine hydroxylase (TH) levels in the rat brain. **(A)** Representative 40μm brain slice at 20X magnification demonstrating delimitation for nucleus accumbens core and shell (NAcC and NAcS), prelimbic cortex (PrLC) and infralimbic cortex (ILC) used for the analysis. Section is immunohistochemically stained for TH (Green) and Hoechst (Blue). **(B)** Detailed TH staining in NAcC, NAcS, ILC and PrLC regions of representative saline- and ACTH-treated animals. **(C)** TH immunofluorescence is presented in arbitrary intensity units. Bars represent means and error bars represent standard error. *=p<0.05 and ns=Non-significant according to unpaired t-test.

As demonstrated in Figure 3, number and volume of IBA1+ cells did not differ between saline and ACTH treatment groups in any of the brain regions analysed according to unpaired t-tests (p>0.05).

**Figure 3:**
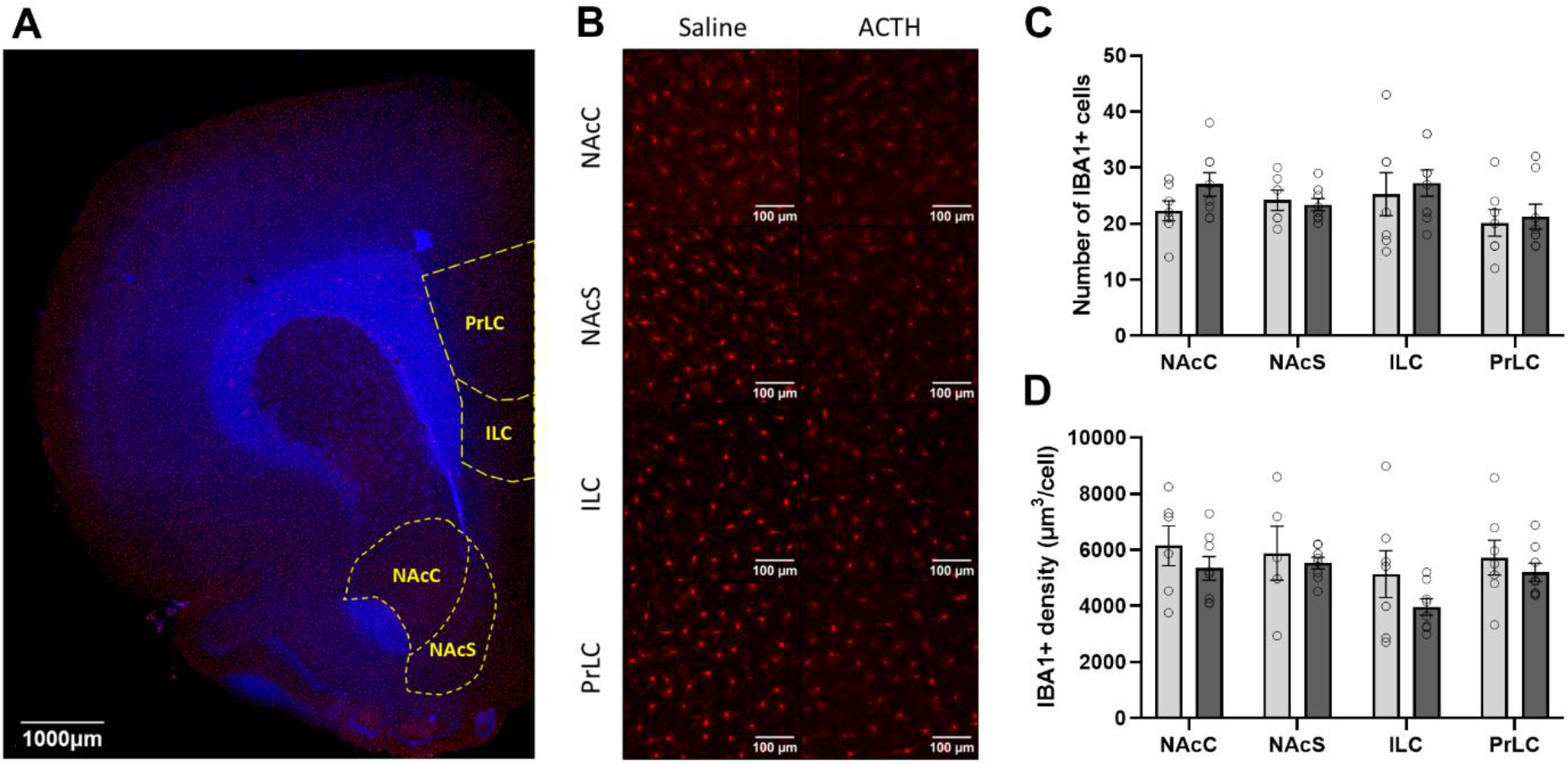
Effects of chronic adrenocorticotrophic hormone (ACTH) or saline treatment on microglia cell levels in the rat brain. **(A)** Representative 40μm brain slice at 20X magnification demonstrating delimitation for nucleus accumbens core and shell (NAcC and NAcS), prelimbic cortex (PrLC) and infralimbic cortex (ILC) used for the analysis. Section is immunohistochemically stained for TH (Green) and Hoechst (Blue). **(B)** Detailed TH staining in NAcC, NAcS, ILC and PrLC regions of representative saline- and ACTH-treated animals. **(C)** TH immunofluorescence is presented in arbitrary intensity units. Bars represent means and error bars represent standard error. No statistical significance was found according to unpaired t-test.

Unpaired t-tests between experimental groups revealed chronic ACTH administration significantly increased IL-6 levels in serum (t=2.364, difference between means ± SEM = 16.40 ± 6.939, p=0.0407) and brain (t=2.211, difference between means ± SEM = 0.01206 ± 0.005456, p=0.0456) compared to control treatment (Fig 4A and 4B). There were no significant differences in serum CRP and PBMC NF-κB levels between experimental groups (Fig 4C and 4D).

**Figure 4:**
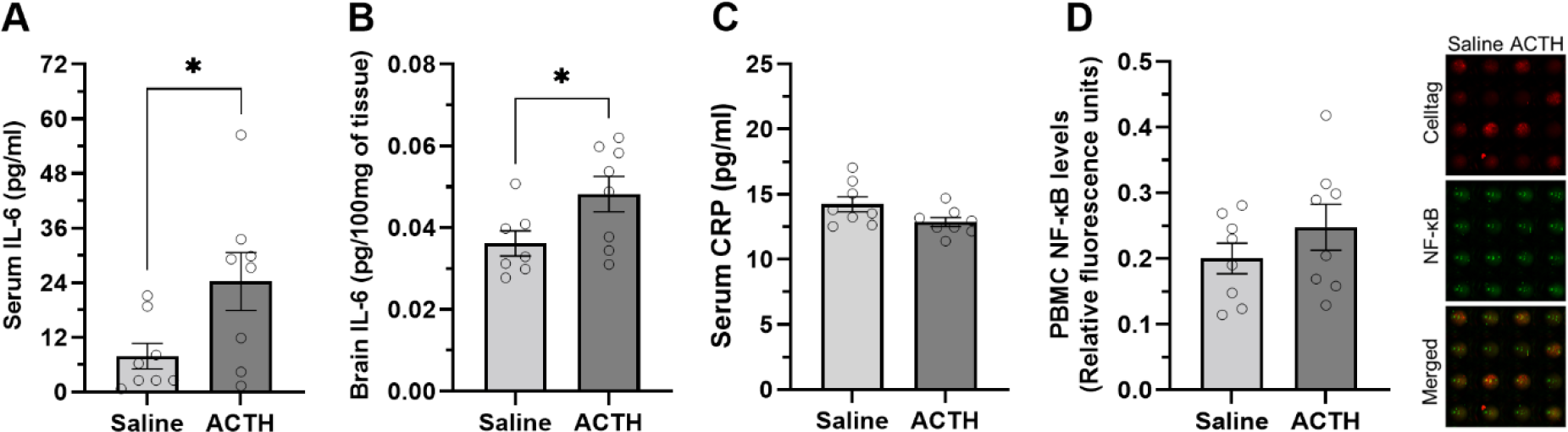
Effects of chronic adrenocorticotrophic hormone (ACTH) or saline treatment on markers of peripheral inflammation. **(A)** Serum interleukin (IL)-6 levels; **(B)** brain IL-6 levels; **(C)** serum C-reactive protein (CRP) levels; and **(D)** peripheral blood mononuclear cells (PBMC) nuclear factor (NF)-κB (B) levels. Bars represent mean and error bars represent standard error, *p<0.05 and ns = non-significant according to unpaired t-test. Image depicts representative In-Cell Western assay for detecting NF-κB in PBMCs, including NF-κB, Cell Tag 700, and merged channels.

PBMCs from ACTH-treated animals showed decreased glucose uptake compared to the control group after insulin stimulation (ACTH treatment: F(1.28)=11.43, p=0.0021; Insulin stimulation: F(1.28)=0.1361, p=0.7150; Interaction: F(1.28)=1.766, p=0.1946) (Fig 5A). When corrected by baseline levels, the ACTH group showed a greater decrease compared to saline-treated animals (t=2.736, difference between means ± SEM = -34.29 ± 12.54, p=0.0161) (Figure 5B).

**Figure 5:**
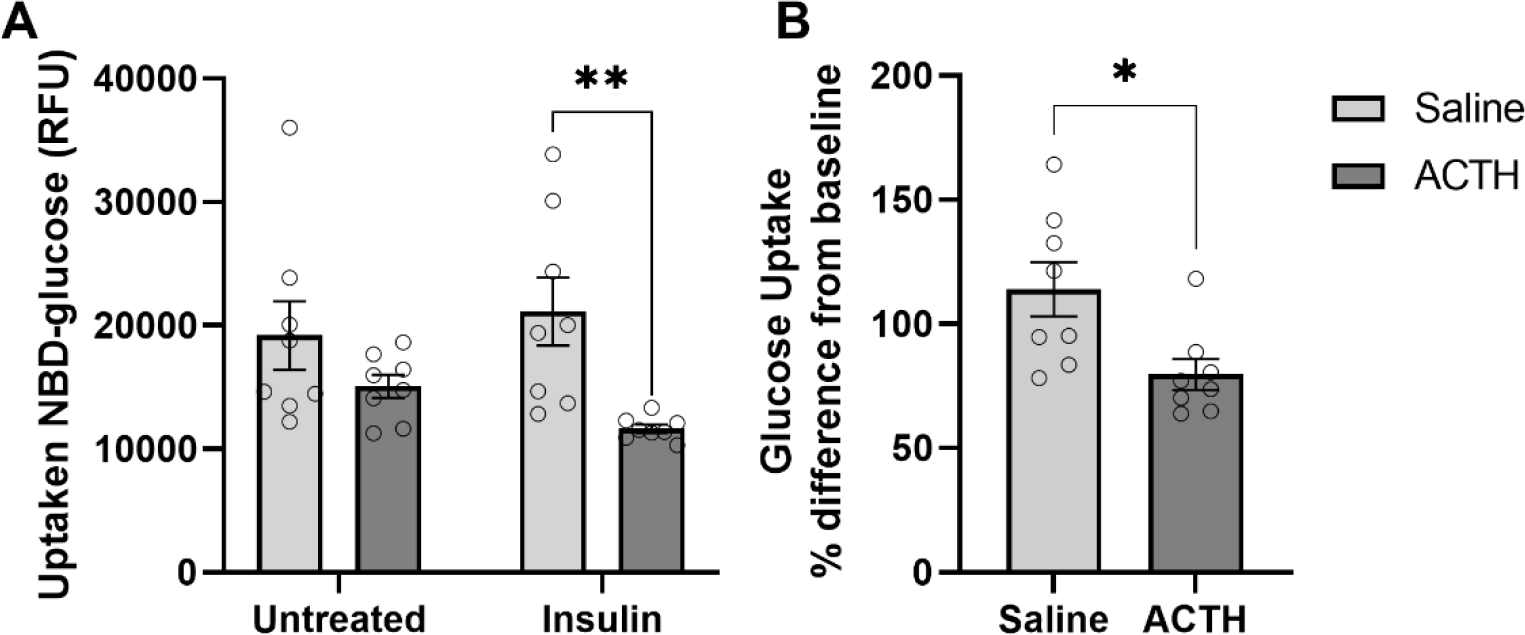
Effects of chronic adrenocorticotrophic hormone (ACTH) or saline treatment on peripheral blood mononuclear cells (PBMC) glucose uptake. **(A)** Intracellular levels of 2-NBD glucose in untreated and insulin-stimulated PBMCs are presented in Relative fluorescence Units (RFU). Bars represent mean and error bars represent standard error, **=p<0.01 according to 2way ANOVA. **(B)** Glucose uptake percentage difference from baseline in ACTH- and saline-treated animals. Bars represent mean and error bars represent standard error, *=p<0.05 according to unpaired t-test.

Metabolomics analysis detected a decrease in the nucleotides adenosine monophosphate (AMP) (U=11, p=0.0281), uridine monophosphate (UMP) (t=2.131, difference between means ± SEM = 863.7 ± 405.3, p=0.05), uridine diphosphate Nacetylglucosamine (UDPNAG) (U=9, p=0.0148), and nicotinamide adenine dinucleotide (oxidised) (NAD) (U=11, p=0.0281) in ACTH-treated animals compared to saline (Figure 6 A, B, C and D). Although not statistically significant, a trend to decrease was also observed in the pentose phosphate pathway, as observed by the markers ribulose-5-phosphate (RL5P) (t=1.414, p=0.0896), xylulose-5-phosphate (X5P) (t=1.450, p=0.0846), ribose-5-phosphate (R5P) (t=1.542, p=0.0726) and erytrose-4-phosphate (E4P) (t=0.3267, p=0.3744) (Figure 6 E, F, G and H). No differences were observed in the metabolites from glycolysis, TCA cycle and others, as demonstrated in supplementary figure 1.

**Figure 6:**
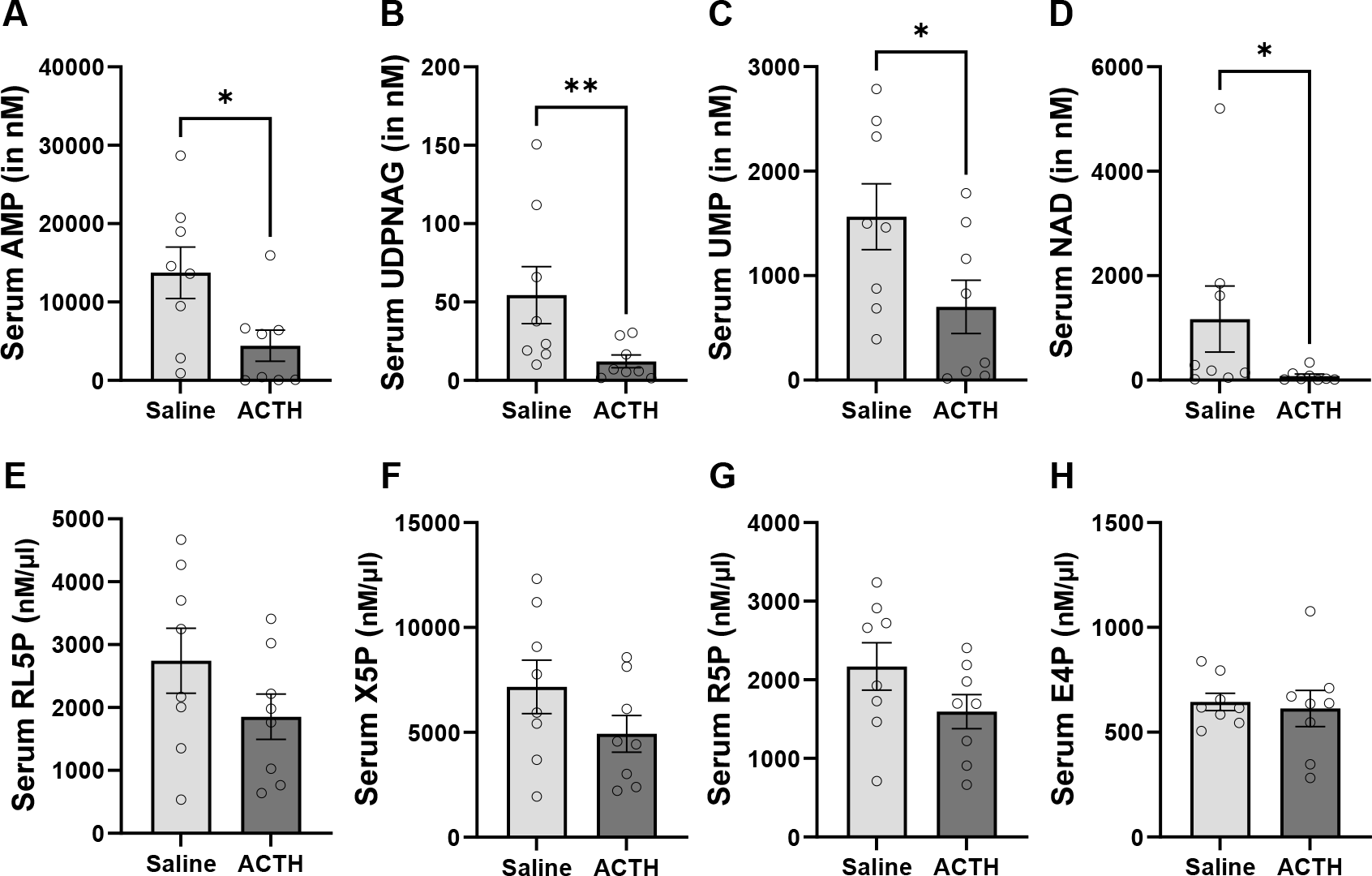
Effects of chronic adrenocorticotrophic hormone (ACTH) or saline treatment on serum glucose metabolism markers concentration. **(A)** Adenosine monophosphate (AMP) levels; **(B)** uridine diphosphate Nacetylglucosamine (UDPNAG) levels; **(C)** uridine monophosphate (UMP) levels; **(D)** nicotinamide adenine dinucleotide (oxidised) (NAD) levels; **(E)** ribulose-5-phosphate (RL5P) levels; **(F)** xylulose-5-phosphate (X5P) levels; **(G)** ribose-5-phosphate (R5P); and **(H)** erythrose-5-phosphate (E4P) levels. Bars represent mean and error bars represent standard error, *=p<0.05; **=p<0.01 according to unpaired t-test.

Simple linear regression revealed significant positive correlations between changes in lever pressing and rewards in the ERCT with PBMC insulin-stimulated glucose uptake (Figure 7A). Levels of TH in the NAcC, ILC and PrLC were negatively correlated to pentose phosphate pathway metabolites RL5P, X5P, R5P, and to the nucleotide guanosine monophosphate (GMP) (Figure 7B, 7C and 7D). The volume of IBA1+ cells was also correlated to local levels of TH in the ILC and PrLC (Figure 7E). IL-6 serum levels were also negatively correlated to RL5P, X5P, R5P, AMP and UMP serum levels (Figure 7F), and ILC TH levels (Figure 7G). All statistical data for significant linear regressions are presented colour-coded in figure 7.

**Figure 7:**
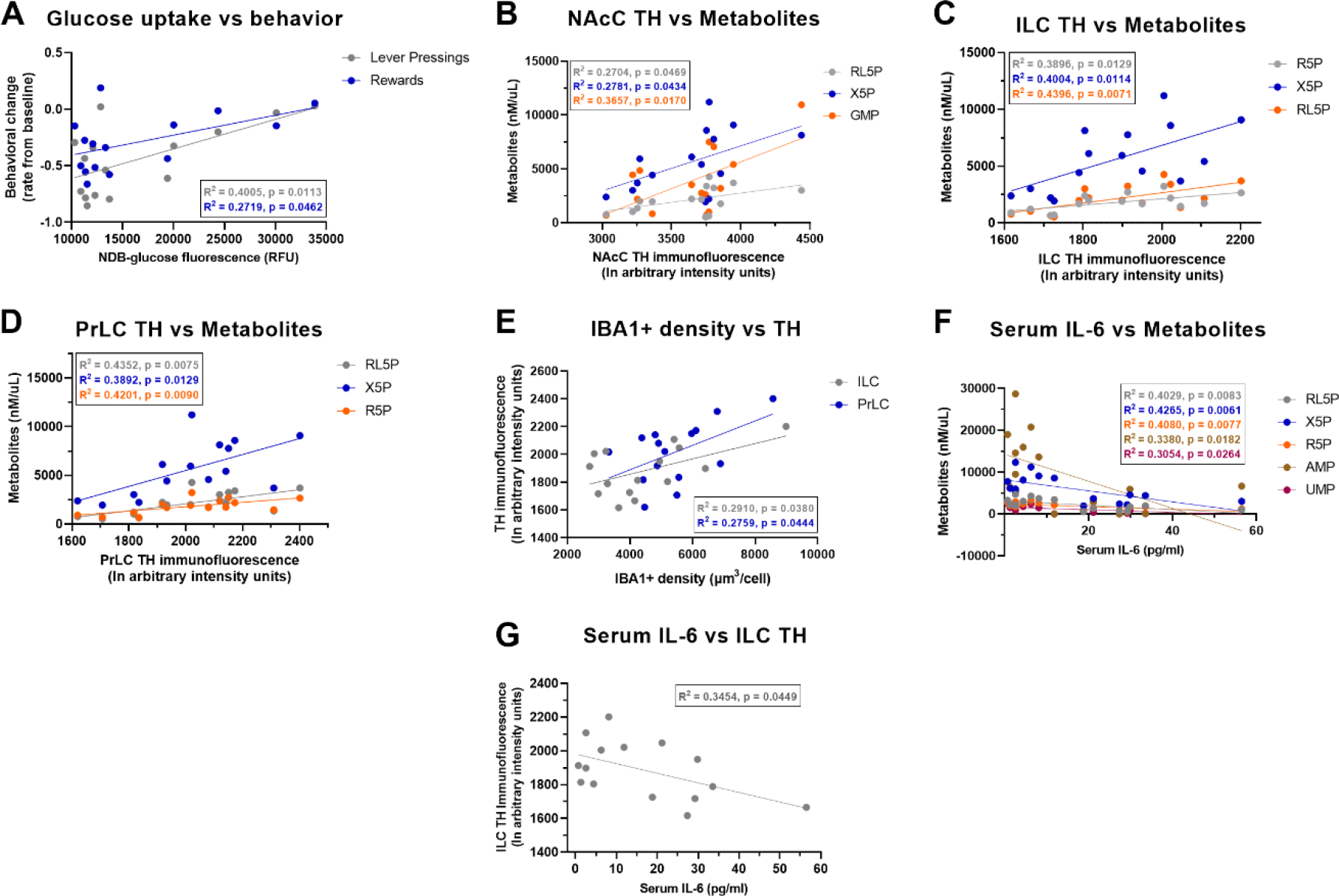
Correlations between multiple behavioural, dopaminergic, metabolic and inflammatory markers. Scatterplots of the linear regression models of **(A)** insulin-stimulated intracellular NDB-glucose fluorescence and changes in lever pressing (in grey) and rewards (in blue) in the effort-related choice task; **(B)** nucleus accumbens core (NacC) tyrosine hydroxylase (TH) levels and serum concentration of D-ribulose 5-phosphate (RL5P) (in grey), xylulose 5-phosphate (X5P) (in blue) and guanosine monophosphate (GMP) (in orange); **(C)** infralimbic cortex (ILC) TH levels and serum concentration of ribose 5-phosphate (R5P) (in grey), X5) (in blue) and RL5P (in orange);**(D)**prelimbic cortex (PrLC) TH levels and serum concentration of RL5P (in grey), X5P (in blue) and R5P (in orange);**(E)**IBA1+ cell density and TH levels in ILC (in grey) and PrLC (in blue); **(F)** serum interleukin-6 (IL-6) levels and serum concentration of RL5P (in grey), X5P (in blue), R5P (in orange), adenosine monophosphate (AMP) (in brown) and uridine monophosphate (in purple) and; **(G)** serum IL-6 levels and ILC TH levels. Simple linear regression R^2^ and p-values are colour-coded for each parameter.

## DISCUSSION

Chronic ACTH injections have been long used as a model of antidepressant resistance, blocking the effects of classical antidepressants such imipramine [23], fluoxetine and bupropion [26], but responding to TRD treatments like ketamine [22] and deep brain stimulation (DBS) [27] in the forced swim test (FST). Although the FST is a good tool to investigate the antidepressant potential of compounds, it was not designed to assess depressive-like behaviours [28]; thus, more translational depressive-like domains, such as anhedonia, are better suited for the full characterisation of a TRD model [29]. In the present study, ACTH treatment was able to reduce lever pressing and rewards consumed in the ERCT when compared to the control group, indicating that chronic injections of ACTH can induce an anhedonia-like phenotype in rats, thereby fulfilling face-validity gaps for a relevant animal model of treatment-resistant depression.

Anhedonia is a DA-dependent behaviour and a core feature for depression disorders. In both animal models and depressive patients, the most relevant biological mechanism of anhedonia is the ventral striatal hypoactivation, but dysfunctions in prefrontal cortex also play an important role in abnormal reward processing and impaired reward-related decision making [30]. In addition to changes in behavioural phenotype, our results demonstrated that chronic ACTH injection decreased TH levels in NAcS and ILC. TH is a key enzyme involved in the process of catecholamine synthesis, regulating DA production and its levels in the brain [31], suggesting that the anhedonia-like phenotype induced by ACTH in the present study might reflect an impaired DA synthesis and consequent altered signalling in these core brain regions of reward processing.

Bupropion is an aminoketone antidepressant which acts by weakly inhibiting reuptake of DA and norepinephrine, with efficacy similar to classical antidepressants [32]. Since bupropion acts through the modulation of DA systems, the effects of acute bupropion treatment were tested in the present study; however, neither 10mg nor 20mg of bupropion were able to rescue the anhedonia phenotype induced by ACTH. In fact, a recent review has demonstrated that bupropion monotherapy does not present efficacy in TRD treatment, and its efficacy when used in augmentation and association with lithium [33]. A potential mechanism underlying bupropion resistance in this model can be explained by the impaired DA synthesis state induced by ACTH. A weak inhibition of DA reuptake might not be enough to overcome low DA availability. Therapies targeting DA synthesis might be more efficient in this scenario, as demonstrated in non-human primates where levodopa (DA precursor) reverses cytokine-induced reductions in striatal DA release and effort-based sucrose consumption [34, 35].

Several studies have suggested that peripheral inflammation is an important feature of depression pathophysiology, although its mechanisms are not fully understood [36]. Chronic stress and HPA axis disruption are known to increase circulating levels of microbe-associated molecular patterns (MAMPs) and danger-associated molecular patterns (DAMPs) [37, 38], which can activate peripheral immune cells leading to a so-called chronic low-grade inflammation state [39]. Activation of peripheral immune cells can induce two different phenotypes – anti-inflammatory or pro-inflammatory states – depending on stimulator patterns [40]. The classic pro-inflammatory phenotype is characterised by activation of the NF-κB pathway, and increased production and release of the proinflammatory cytokines TNF, IL-1 and IL-6 [41, 42]. Although the increase in PBMCs NF-κB levels were not significant in this study, ACTH-treated animals had increased serum levels of IL-6 when compared to saline-treated controls, demonstrating an increase in proinflammatory signalling in the present model.

Peripheral inflammation can affect the brain in many ways, either inducing neuroinflammation, directly altering neurotransmitter systems or both [43]. Proinflammatory cytokines can directly enter the brain through blood-brain barrier leakage or transport protein channels [44, 45], stimulate release of inflammatory mediators by endothelial cells and perivascular macrophage in the brain [46], or induce peripheral monocyte infiltration [47]. IL-6 plays a major role in this peripheral-brain communication, and it seems to be involved in both normal and abnormal physiological states [46, 48, 49]. In fact, we were able to demonstrate increased levels of IL-6 in serum and brain of animals treated with ACTH compared to controls, suggesting its role as a periphery-brain signalling molecule in this context. Although serum and central levels of IL-6 were not directly correlated in our study, it corroborates with a recent meta-analysis study demonstrating the lack of general blood-CSF correlation for interleukins [50], suggesting indirect mechanisms of communication and/or the involvement of other mediators in this process.

DAergic systems are sensitive to peripheral inflammation, particularly increases in IL-6 levels [51]. Although this is not fully understood, the potential mechanisms include changes in DA synthesis, packing, transport, reuptake, and receptors [52]. Mice presenting depressive-like phenotype after chronic social stress demonstrated increased BBB permeability and trafficking of IL-6 from the periphery to the NAc, a major DAergic hub in the brain [53]. Interestingly, a clinical study also demonstrated that peripheral levels of IL-6 are correlated to individual differences on NAc prediction error signals measured with fMRI after acute stress [54]. A clinical study has demonstrated a correlation between IL-6 levels and increase in phenylalanine concentration (a precursor of tyrosine) in cancer patients [49]. IL-6 treatment in animals was also shown to decrease tetrahydrobiopterin (BH4) in presynaptic neurons, a TH enzyme cofactor [50]. Corroborating with these observations, the present study demonstrates a negative correlation between serum IL-6 levels and TH levels in the ILC. Also, although no group differences were observed, increased local IBA1+ cell density, a marker of ramified physiological microglia, was positively correlated to TH levels in the ILC and PrLC. Altogether, this data suggests that peripheral inflammatory markers, IL-6 especially, can affect central DA metabolism by decreasing availability of enzymes and cofactors involved in its synthesis, leading to depressive-like symptoms.

These results highlight the link between peripheral and neuroinflammation to DA dysregulation and consequent disruption of reward-related behaviours in the present model. In the context of low-grade inflammation, this crosstalk functions to signal the brain to support the metabolic demands and perceived threats to body integrity and is able to modulate entire neurotransmitter systems and complex behavioural responses [55]. In fact, our results demonstrate that ACTH treatment decreases peripheral insulin-stimulated glucose uptake. ACTH is the main secretory hormone of glucocorticoids, which can lead to insulin resistance directly [56] or through different pathways, such fatty acid accumulation [57], desensitisation of insulin receptors and decreased insulin-mediated expression of glucose transporter in the membrane [58]. Abnormal levels of ACTH are associated with insulin resistance in many patients with metabolic conditions [59], demonstrating the clinical relevance of HPA axis disruption in metabolic changes. In the present study, metabolomics analysis shown decreased nucleotide synthesis and, although not significant, a trend for impaired pentose phosphate pathway activity (PPP) in the serum of ACTH-treated animals. The glycolysis and PPP are tightly connected, as the glucose uptaken by the cell is phosphorylated and converted to glucose 6-phosphate (G6P). G6P can then be metabolised either by the glycolytic pathway, generating pyruvate and lactate, or by PPP to produce NADPH and nucleotides [60]. The glucose flux between the PPP and glycolysis is regulated by several mechanisms, and when the glycolytic pathway is inhibited, glucose re-flux towards the PPP is increased. This shift is dynamic and can adapt according to energy availability/demand in a feedback loop manner [60]. Thus, our results suggest that ACTH may induce insulin resistance and decreased glucose uptake in peripheral cells. Reduced intracellular glucose is shifted towards glycolytic pathway to support energy demands, resulting in reduced PPP activity and nucleotide synthesis.

Correlations between serum IL-6 levels and metabolic markers in the present study indicate a metabolic shift towards glycolysis, suggesting an adaptative strategy to supply energy for peripheral immune cell activation and increased production of inflammatory mediators. Interestingly, levels of TH were positively correlated to energy metabolism markers and microglial cell density, and negatively correlated to serum IL-6 levels in a region-dependent manner, suggesting IL-6 is the mediator between peripheral metabolic/inflammatory demands and modulation of DA-dependent behaviours.

Some limitations must be acknowledged when interpreting the results from the present study. Bupropion results must be interpreted carefully since no response was observed in saline-treated animals, contrasting with results from a similar study by Randall and colleagues [61] demonstrating that 20mg (but not 10 mg) of bupropion was able to increase rats’ performance in the same behavioural paradigm. However, the discrepancy in the results might be explained by differences in experimental design. In the present study, control animals received chronic intraperitoneal injections of saline every day for 24 days, whilst in the Randall study, animals received only acute injections before the test session. Chronic stress of daily injections by itself might affect bupropion response, and a higher dose is necessary to induce behavioural changes in the control group.

Interestingly, although increased glycolytic activity is the hallmark for immune cell activation, insulin resistance is a feature of the anti-inflammatory phenotype M2 in peripheral immune cells [62]. Since the focus of this study is the role of IL-6 as a periphery-brain crosstalk mediator, no anti-inflammatory interleukin level was assessed. Further studies evaluating the balance between pro- and anti-inflammatory interleukins will provide a better understanding of general inflammatory state induced by chronic ACTH. It is important to note that together with glycolysis, increased PPP activity is often observed as part of a metabolic shift induced by chronic low-grade inflammation, and its bioproducts are important for FA synthesis [62]; thus, the decrease in serum PPP pathway metabolites observed in the present study seems controversial. However, this discrepancy might be explained by the insulin resistance induced by ACTH, which might decrease intracellular glucose availability, acting as a limiting factor for proper metabolic response. In this scenario, the glucose would preferentially flux through glycolytic pathway to support ATP production, decreasing PPP activity. It is also noteworthy that in the brain PPP modulates chronic neuroinflammation and DAergic neurodegeneration [63] and is inhibited by antidepressants [64]. In the present study we only analyse PPP metabolites in serum, which can have a different profile from the brain. Future studies assessing cerebral PPP activity under this context are necessary to demonstrate how this pathway is differentially modulated.

## Conclusion and future directions

The present study demonstrated that chronic injections of ACTH can induce anhedonia-like phenotype in rats, a translational symptom domain of clinical depression. This anhedonia phenotype seems to be resistant to bupropion, a DA modulator antidepressant, highlighting the classical antidepressant resistance nature of this model. The anhedonia-like phenotype induced by ACTH seems to be constructed upon peripherical metabolic changes and increased inflammation triggering neuroimmune activation and impairment of DA metabolism. These results reinforce not only its validities of a TRD model, but also offer an important tool to investigate the cross talk between peripheral and central alterations.

Despite its potential applications, several aspects need to be further characterised in this model, such as a better predictive profile including different doses of bupropion, but also other therapies efficient for TRD treatment (such as ketamine and DBS). A better description of inflammatory profile and metabolic alterations are also needed in order to understand in detail how peripheral state can modulate motivated behaviours.

## Acknowledgments

The authors gratefully acknowledge Dr Clarissa Yates, Dr Nathanael Yates and the QBI Advanced Microscopy and Histology facilities for their support and assistance in this work.

## Notes

### Competing Interest Statement

The authors have declared no competing interest.

### Summary of Updates

Inclusion of co-author responsible for metabolomics analysis and data interpretation

## References

1. Baik, J.-H., Stress and the dopaminergic reward system. Experimental & Molecular Medicine, 2020. 52(12): p. 1879–1890.

2. Bromberg-Martin, E.S., M. Matsumoto, and O. Hikosaka, Dopamine in motivational control: rewarding, aversive, and alerting. Neuron, 2010. 68(5): p. 815–34.

3. Husain, M. and J.P. Roiser, Neuroscience of apathy and anhedonia: a transdiagnostic approach. Nature Reviews Neuroscience, 2018. 19(8): p. 470–484.

4. Ferenczi, E.A., et al., Prefrontal cortical regulation of brainwide circuit dynamics and reward-related behavior. Science, 2016. 351(6268): p. aac9698.

5. Stringaris, A., et al., The Brain’s Response to Reward Anticipation and Depression in Adolescence: Dimensionality, Specificity, and Longitudinal Predictions in a Community-Based Sample. Am J Psychiatry, 2015. 172(12): p. 1215–23.

6. American-Psychiatric-Association, Diagnostic and statistical manual of mental disorders : DSM-5, ed. A. American Psychiatric and D.S.M.T.F. American Psychiatric Association. 2013, Arlington, VA: American Psychiatric Association.

7. Cooper, J.A., A.R. Arulpragasam, and M.T. Treadway, Anhedonia in depression: biological mechanisms and computational models. Current Opinion in Behavioral Sciences, 2018. 22: p. 128–135.

8. Miller, A.H. and C.L. Raison, The role of inflammation in depression: from evolutionary imperative to modern treatment target. Nature Reviews Immunology, 2016. 16(1): p. 22–34.

9. Gaynes, B.N., et al., The STAR*D study: treating depression in the real world. Cleve Clin J Med, 2008. 75(1): p. 57–66.

10. Ruberto, V.L., M.K. Jha, and J.W. Murrough, Pharmacological Treatments for Patients with Treatment-Resistant Depression. Pharmaceuticals (Basel), 2020. 13(6).

11. Dowlati, Y., et al., A meta-analysis of cytokines in major depression. Biol Psychiatry, 2010. 67(5): p. 446–57.

12. Hiles, S.A., et al., A meta-analysis of differences in IL-6 and IL-10 between people with and without depression: exploring the causes of heterogeneity. Brain Behav Immun, 2012. 26(7): p. 1180–8.

13. Tang, W., et al., Inflammatory cytokines, complement factor H and anhedonia in drug-naïve major depressive disorder. Brain Behav Immun, 2021. 95: p. 238–244.

14. Felger, J.C., et al., Inflammation is associated with decreased functional connectivity within corticostriatal reward circuitry in depression. Molecular Psychiatry, 2016. 21(10): p. 1358–1365.

15. McMakin, D.L., et al., Anhedonia predicts poorer recovery among youth with selective serotonin reuptake inhibitor treatment-resistant depression. J Am Acad Child Adolesc Psychiatry, 2012. 51(4): p. 404–11.

16. Krepel, N., et al., Can psychological features predict antidepressant response to rTMS? A Discovery-Replication approach. Psychol Med, 2020. 50(2): p. 264–272.

17. Bonanni, L., et al., Can Anhedonia Be Considered a Suicide Risk Factor? A Review of the Literature. Medicina (Kaunas), 2019. 55(8).

18. Thomas, R.K., et al., Rapid effectiveness of intravenous ketamine for ultraresistant depression in a clinical setting and evidence for baseline anhedonia and bipolarity as clinical predictors of effectiveness. Journal of Psychopharmacology, 2018. 32(10): p. 1110–1117.

19. Uliana, D.L., et al., Using animal models for the studies of schizophrenia and depression: The value of translational models for treatment and prevention. Front Behav Neurosci, 2022. 16: p. 935320.

20. Insel, T., et al., Research domain criteria (RDoC): toward a new classification framework for research on mental disorders. Am J Psychiatry, 2010. 167(7): p. 748–51.

21. Anderzhanova, E., T. Kirmeier, and C.T. Wotjak, Animal models in psychiatric research: The RDoC system as a new framework for endophenotype-oriented translational neuroscience. Neurobiol Stress, 2017. 7: p. 47–56.

22. Walker, A.J., et al., Peripheral proinflammatory markers associated with ketamine response in a preclinical model of antidepressant-resistance. Behav Brain Res, 2015. 293: p. 198–202.

23. Walker, A.J., et al., Chronic adrenocorticotrophic hormone treatment alters tricyclic antidepressant efficacy and prefrontal monoamine tissue levels. Behav Brain Res, 2013. 242: p. 76–83.

24. Randall, P.A., et al., Dopaminergic modulation of effort-related choice behavior as assessed by a progressive ratio chow feeding choice task: pharmacological studies and the role of individual differences. PLoS One, 2012. 7(10): p. e47934.

25. Schindelin, J., et al., Fiji: an open-source platform for biological-image analysis. Nature Methods, 2012. 9(7): p. 676–682.

26. Srikumar, B.N., et al., Characterization of the adrenocorticotrophic hormone - induced mouse model of resistance to antidepressant drug treatment. Pharmacol Biochem Behav, 2017. 161: p. 53–61.

27. Kale, R.P., et al., Mood Regulatory Actions of Active and Sham Nucleus Accumbens Deep Brain Stimulation in Antidepressant Resistant Rats. Front Hum Neurosci, 2021. 15: p. 644921.

28. Commons, K.G., et al., The Rodent Forced Swim Test Measures Stress-Coping Strategy, Not Depression-like Behavior. ACS Chem Neurosci, 2017. 8(5): p. 955–960.

29. Planchez, B., A. Surget, and C. Belzung, Animal models of major depression: drawbacks and challenges. J Neural Transm (Vienna), 2019. 126(11): p. 1383–1408.

30. Serretti, A., Anhedonia and Depressive Disorders. Clin Psychopharmacol Neurosci, 2023. 21(3): p. 401–409.

31. Daubner, S.C., T. Le, and S. Wang, Tyrosine hydroxylase and regulation of dopamine synthesis. Arch Biochem Biophys, 2011. 508(1): p. 1–12.

32. Stahl, S.M., et al., A Review of the Neuropharmacology of Bupropion, a Dual Norepinephrine and Dopamine Reuptake Inhibitor. Prim Care Companion J Clin Psychiatry, 2004. 6(4): p. 159–166.

33. Tran, K., S.C. McGill, and J. Horton, CADTH Health Technology Review, in Bupropion for Treatment-Resistant Depression. 2021, Canadian Agency for Drugs and Technologies in Health. Copyright © 2021 Canadian Agency for Drugs and Technologies in Health.: Ottawa (ON).

34. Felger, J.C., C.R. Hernandez, and A.H. Miller, Levodopa reverses cytokine-induced reductions in striatal dopamine release. Int J Neuropsychopharmacol, 2015. 18(4).

35. Felger, J.C., et al., Chronic interferon-α decreases dopamine 2 receptor binding and striatal dopamine release in association with anhedonia-like behavior in nonhuman primates. Neuropsychopharmacology, 2013. 38(11): p. 2179–87.

36. Suneson, K., et al., An inflamed subtype of difficult-to-treat depression. Prog Neuropsychopharmacol Biol Psychiatry, 2023. 125: p. 110763.

37. Fleshner, M., M. Frank, and S.F. Maier, Danger Signals and Inflammasomes: Stress-Evoked Sterile Inflammation in Mood Disorders. Neuropsychopharmacology, 2017. 42(1): p. 36–45.

38. Maslanik, T., et al., Commensal bacteria and MAMPs are necessary for stress-induced increases in IL-1β and IL-18 but not IL-6, IL-10 or MCP-1. PLoS One, 2012. 7(12): p. e50636.

39. Sen, Z.D., et al., Linking atypical depression and insulin resistance-related disorders via low-grade chronic inflammation: Integrating the phenotypic, molecular and neuroanatomical dimensions. Brain Behav Immun, 2021. 93: p. 335–352.

40. Ley, K., M1 Means Kill; M2 Means Heal. J Immunol, 2017. 199(7): p. 2191–2193.

41. Dorrington, M.G. and I.D.C. Fraser, NF-κB Signaling in Macrophages: Dynamics, Crosstalk, and Signal Integration. Front Immunol, 2019. 10: p. 705.

42. Margraf, A. and M. Perretti, Immune Cell Plasticity in Inflammation: Insights into Description and Regulation of Immune Cell Phenotypes. Cells, 2022. 11(11).

43. Sun, Y., Y. Koyama, and S. Shimada, Inflammation From Peripheral Organs to the Brain: How Does Systemic Inflammation Cause Neuroinflammation? Front Aging Neurosci, 2022. 14: p. 903455.

44. Obermeier, B., R. Daneman, and R.M. Ransohoff, Development, maintenance and disruption of the blood-brain barrier. Nat Med, 2013. 19(12): p. 1584–96.

45. Banks, W.A., Blood-brain barrier transport of cytokines: a mechanism for neuropathology. Curr Pharm Des, 2005. 11(8): p. 973–84.

46. Dantzer, R., et al., From inflammation to sickness and depression: when the immune system subjugates the brain. Nat Rev Neurosci, 2008. 9(1): p. 46–56.

47. D’Mello, C., T. Le, and M.G. Swain, Cerebral microglia recruit monocytes into the brain in response to tumor necrosis factoralpha signaling during peripheral organ inflammation. J Neurosci, 2009. 29(7): p. 2089–102.

48. Erta, M., A. Quintana, and J. Hidalgo, Interleukin-6, a major cytokine in the central nervous system. Int J Biol Sci, 2012. 8(9): p. 1254–66.

49. Garner, K.M., et al., Microglia priming by interleukin-6 signaling is enhanced in aged mice. J Neuroimmunol, 2018. 324: p. 90–99.

50. Gigase, F.A.J., et al., The association between inflammatory markers in blood and cerebrospinal fluid: a systematic review and meta-analysis. Mol Psychiatry, 2023. 28(4): p. 1502–1515.

51. Jones, B.D.M., et al., Associations between peripheral inflammation and clinical phenotypes of bipolar depression in a lower-middle income country. CNS Spectr, 2023: p. 1–9.

52. Felger, J.C. and M.T. Treadway, Inflammation Effects on Motivation and Motor Activity: Role of Dopamine. Neuropsychopharmacology, 2017. 42(1): p. 216–241.

53. Menard, C., et al., Social stress induces neurovascular pathology promoting depression. Nat Neurosci, 2017. 20(12): p. 1752–1760.

54. Treadway, M.T., et al., Association Between Interleukin-6 and Striatal Prediction-Error Signals Following Acute Stress in Healthy Female Participants. Biol Psychiatry, 2017. 82(8): p. 570–577.

55. Treadway, M.T., J.A. Cooper, and A.H. Miller, Can’t or Won’t? Immunometabolic Constraints on Dopaminergic Drive. Trends Cogn Sci, 2019. 23(5): p. 435–448.

56. Iwen, K.A., et al., Melanocortin crosstalk with adipose functions: ACTH directly induces insulin resistance, promotes a pro-inflammatory adipokine profile and stimulates UCP-1 in adipocytes. J Endocrinol, 2008. 196(3): p. 465–72.

57. Geer, E.B., J. Islam, and C. Buettner, Mechanisms of glucocorticoid-induced insulin resistance: focus on adipose tissue function and lipid metabolism. Endocrinol Metab Clin North Am, 2014. 43(1): p. 75–102.

58. Leonard, B.E. and G. Wegener, Inflammation, insulin resistance and neuroprogression in depression. Acta Neuropsychiatr, 2020. 32(1): p. 1–9.

59. Guan, W., et al., Endocrine characteristics and risk factors of type 2 diabetes complicated with gastrointestinal autonomic neuropathy: A single-center retrospective study. Medicine (Baltimore), 2023. 102(15): p. e33467.

60. Cho, E.S., et al., The Pentose Phosphate Pathway as a Potential Target for Cancer Therapy. Biomol Ther (Seoul), 2018. 26(1): p. 29–38.

61. Randall, P.A., et al., Bupropion increases selection of high effort activity in rats tested on a progressive ratio/chow feeding choice procedure: implications for treatment of effort-related motivational symptoms. Int J Neuropsychopharmacol, 2014. 18(2).

62. Jin, E.S., et al., Pentose phosphate pathway activity parallels lipogenesis but not antioxidant processes in rat liver. Am J Physiol Endocrinol Metab, 2018. 314(6): p. E543–e551.

63. Tu, D., et al., The pentose phosphate pathway regulates chronic neuroinflammation and dopaminergic neurodegeneration. Journal of Neuroinflammation, 2019. 16(1): p. 255.

64. Özaslan, M.S., et al., Inhibition effects of some antidepressant drugs on pentose phosphate pathway enzymes. Environ Toxicol Pharmacol, 2019. 72: p. 103244.

