## supplemental material for "The role of IL-6 in dopamine dysregulation underlying anhedonia phenotype in rats"

### Supplementary materials

| Pathway | Abbreviation | Compound |
|---|---|---|
| Glycolysis | F16DP | fructose 1,6-bisphosphate |
| | G1P | glucose 1-phosphate |
| | G6P | glucose 6-phosphate |
| | F6P | fructose 6-phosphate |
| | LAC | lactate |
| | PEP | phosphoenolpyruvate |
| | PYR | pyruvate |
| | DHAP | dihydroxyacetone phosphate |
| | G3P | Glycerol-3-phosphate |
| TCA Cycle | ACO | aconitic acid |
| | FUM | fumarate |
| | ISO | isocitrate |
| | CIT | citrate |
| | KGA | α-ketoglutarate |
| | MAL | malate |
| | SUC | succinate |
| Pentose phosphate pathway | E4P | D-erythrose 4-phosphate |
| | R5P | D-ribose 5-phosphate |
| | RL5P | D-ribulose 5-phosphate |
| | X5P | D-xylulose 5-phosphate |
| nucleotides | AMP | adenosine monophosphate |
| | CMP | cytidine monophosphate |
| | GMP | guanosine monophosphate |
| | UDPNAG | uridine diphosphate N-acetylglucosamine |
| | UMP | uridine monophosphate |
| Cofactors | NAD | nicotinamide adenine dinucleotide (oxidised) |
| others | GLYCO | glycolic acid |
| | Cre-PO | creatine phosphate |
| | ADI | adipic acid |

**Supplementary table 1:** Complete list of central carbon metabolism compounds analysed through liquid chromatography–mass spectrometry (LC/MS) in serum samples.

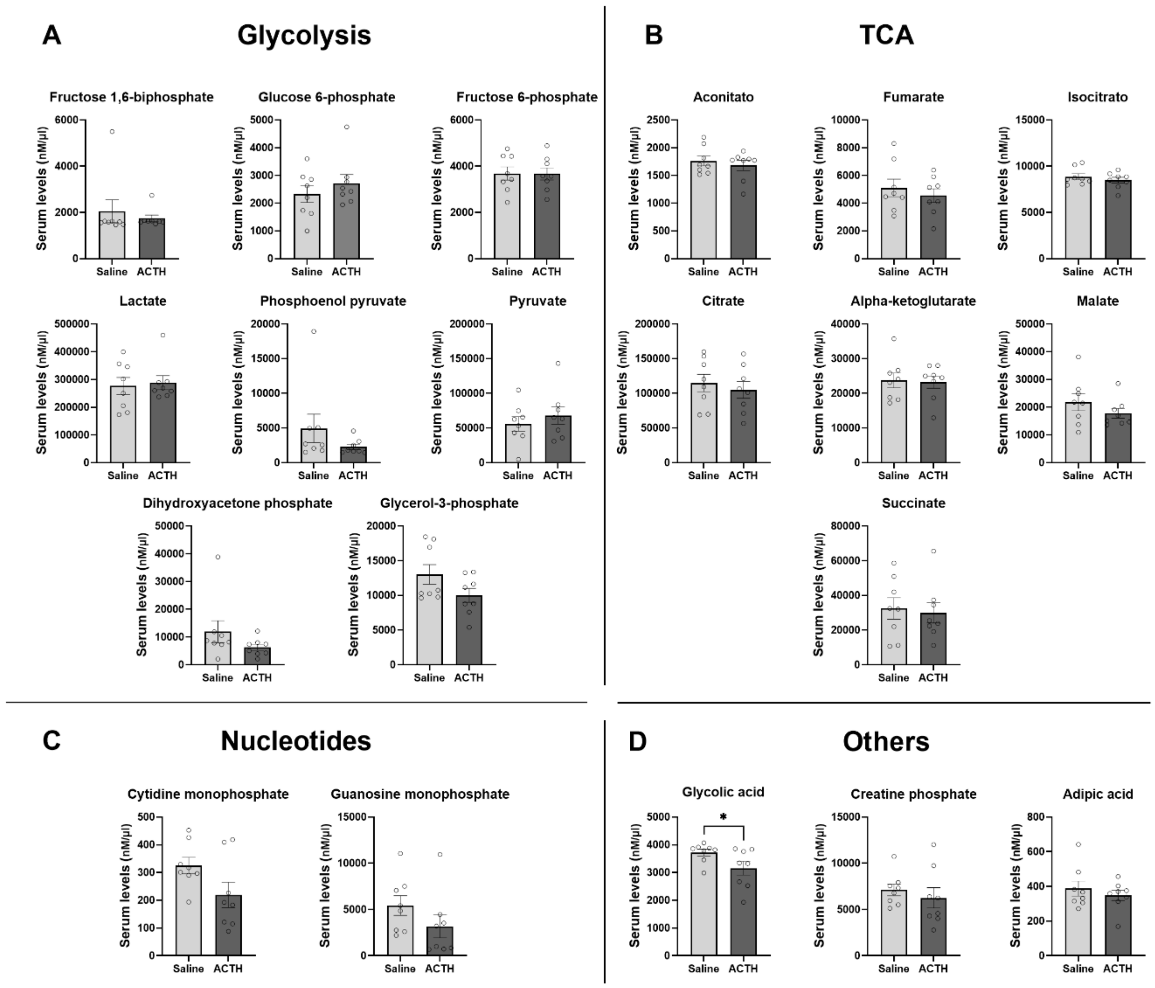


**Supplementary figure 2:** Effects of ACTH and saline administration on serum glucose metabolism marker levels related to glycolysis (A); tricarboxylic acid cycle (TCA) (B); Nucleotides (C); and others (D) metabolites. Bars represent Mean, error bars represent ±SEM. *p<0.05; according to unpaired t-test or Mann-Whitney according to normality test.
